# Excitotoxic glutamate levels cause the secretion of resident endoplasmic reticulum proteins

**DOI:** 10.1101/2023.09.03.556116

**Authors:** Amanda M. Dossat, Kathleen A. Trychta, Lowella V. Fortuno, Christopher T. Richie, Brandon K. Harvey

## Abstract

Dysregulation of synaptic glutamate levels can lead to excitotoxicity such as that observed in stroke, traumatic brain injury, and epilepsy. The role of increased intracellular calcium (Ca^2+^) in the development of excitotoxicity is well established. However, less is known regarding the impact of glutamate on endoplasmic reticulum (ER)-Ca^2+^-mediated processes such as proteostasis. To investigate this, we expressed a secreted ER Ca^2+^ modulated protein (SERCaMP) in primary cortical neurons to monitor exodosis, a phenomenon whereby ER calcium depletion causes the secretion of ER resident proteins that perform essential functions to the ER and the cell. Activation of glutamatergic receptors (GluRs) led to an increase in SERCaMP secretion indicating that normally ER resident proteins are being secreted in a manner consistent with ER Ca^2+^ depletion. Antagonism of ER Ca^2+^ channels attenuated the effects of glutamate and GluR agonists on SERCaMP release. We also demonstrate that endogenous proteins containing an ER retention sequence (ERS) are secreted in response to GluR activation supporting that neuronal activation by glutamate promotes ER exodosis. Ectopic expression of KDEL receptors attenuated the secretion of ERS-containing proteins caused by GluR agonists. Taken together, our data indicate that excessive GluR activation causes disruption of neuronal proteostasis by triggering the secretion of ER resident proteins through ER Ca^2+^ depletion and describes a new facet of excitotoxicity.

**Significance:** During excitotoxicity, the excessive activation of glutamate receptors causes elevated intracellular calcium (Ca^2+^) that promotes cellular dysfunction and death. While the role of cytosolic Ca^2+^ in excitotoxicity has been well-studied, the consequences of changes in endoplasmic reticulum (ER) Ca^2+^ during excitotoxicity remains unclear. The relatively high concentration of calcium in the ER is necessary for ER resident proteins to function prop out essential functions and maintain cellular proteostasis. We show here that excitotoxic conditions destabilize the ER proteome by triggering ER resident protein secretion. Stabilizing ER Ca^2+^ or overexpressing receptors that interact with ER resident proteins can prevent disruption of proteostasis associated with excitotoxicity. The present study provides a new link between excitotoxicity, ER Ca^2+^ homeostasis, and the ER proteome.

## Introduction

Brain injuries such as stroke, traumatic brain injury, and seizure can lead to elevations in extracellular excitatory amino acids (EEAs), primarily glutamate ^1–3^. The excessive synaptic glutamate acts upon postsynaptic ionotropic and metabotropic glutamate receptors to initiate a cascade of downstream intracellular events. More specifically, a surplus of extracellular glutamate leads to dysregulation of intracellular Ca^2+^ and excitotoxicity^4–7^, which contributes to the pathophysiology of several neurodegenerative disorders, including epilepsy and stroke^8^.

The endoplasmic reticulum (ER) is the primary storage site and modulator of intracellular Ca^2+^, in addition to its critical functions associated with the synthesis, modification, and trafficking of lipids and proteins (i.e., proteostasis)^9–11^. Ca^2+^ levels in the ER are estimated to be 5000-fold higher than calcium concentrations found in the cytosol^12^, and this reservoir of intracellular Ca^2+^ supports many cellular functions. We recently discovered that decreased levels of ER Ca^2+^ caused a compositional change in the ER proteome, specifically the secretion of proteins that typically reside in the ER lumen to perform the aforementioned functions of the ER^13^. ER resident proteins contain carboxyl-terminal ER retention sequences (ERS) which interact with KDEL receptors (KDELRs) as part of the KDEL retrieval pathway^14,15^. Under physiological conditions, the KDEL retrieval pathway serves as a mechanism to promote ER localization of proteins containing an ERS tail. ERS-containing proteins (e.g., chaperones, isomerases, etc.) that enter the secretory pathway and travel to the cis-Golgi are recognized by KDELRs^14,16^. The interaction between a carboxy-terminal ERS and KDELR in the Golgi triggers retrograde transport of the proteins back to the ER lumen^17,18^. Under conditions of decreased ER Ca^2+^, the ERS proteins overwhelm the KDELRs and are secreted from the cell in a process referred to as “exodosis”^13^.

A *Gaussia* luciferase (GLuc) with an ERS carboxy tail (GLuc-SERCaMP)^19^, can be used to monitor changes in the ER proteome associated with ER Ca^2+^ depletion. We have demonstrated that reductions in ER Ca^2+^ lead to secretion of ERS proteins and of GLuc-SERCaMP^19,20^. Thus, extracellular levels of GLuc-SERCaMP serve as an indicator of reduced ER Ca^2+^ levels and secretion of resident ER proteins. A previous report from our lab indicated that glutamate caused secretion of GLuc-SERCaMP^19^, and others have reported the ability of excitotoxicity to influence the ER proteome^21,22^. Here, using the bioluminescent reporter of ER Ca^2+^ depletion and ER exodosis (GLuc-SERCaMP)^13,19^, we describe the effects of glutamate on ER exodosis in neurons. Our findings provide evidence that exodosis occurs during excitotoxicity and stabilizing ER Ca^2+^ or increasing KDELR expression mitigates exodosis. This represents a therapeutic target for reducing cellular dysfunction associated with neurological insults and diseases where excessive glutamatergic activity is observed. Future studies examining the role of secreted endogenous ERS proteins in models of excitotoxicity may identify new therapeutic targets for treating elevated extracellular glutamate as observed in conditions such as stroke, traumatic brain injury, or seizure.

## Materials & Methods

### Plasmid Construction

All plasmids were constructed using ligation-independent cloning (In-Fusion, Clontech) into the backbone of pAAV CaMKII iRFP-FLAG (Addgene 149513) after digestion with NheI and AscI restriction enzymes (New England Biolabs) and insert amplification by polymerase chain reaction. All PCR reactions were performed with Q5 polymerase (New England Biolabs) and cleaned up using Nucleospin Gel and PCR spin columns (Macherey-Nagel). In-Fusion reactions were transformed into NEB Stable competent cells (New England Biolabs). Insert-containing clones were subjected to restriction digest analysis and sequence verification prior to subsequent cloning steps or packaging into viral particles. The coding regions for GLuc-no tag and GLuc-SERCaMP were amplified from pLenti6.3 CMV MANF-sigpep-GLuc-MCS and pLenti6.3 CMV MANF-sigpep-GLuc-ASARTDL, respectively^13^ and produced AAV-CaMKIIα-GLuc-no tag (Addgene 149502) and AAV-CaMKIIα-GLuc-SERCaMP (Addgene 149503), respectively . AAV-CamKII-eYFP plasmid was a gift from Dr. Karl Deisseroth.

The coding regions for human KDELRs with Myc-DDK tags, hKDELR1-Myc-DDK and hKDELR2-Myc-DDK, were amplified from previously described plasmids^19^ and produced plasmids pAAV CaMKII KDELR1-Myc-DDK (Addgene# 149514) and pAAV CaMKII KDELR2-Myc-DDK (Addgene# 149515), respectively.

### Viral Packaging

All AAV vectors were produced using triple transfection method as previously described^23^. All vectors were produced using serotype 1 capsid proteins and titered by droplet digital PCR.

### Cell culture

Mixed-sex, rat primary cortical neurons (PCNs) were isolated on embryonic day 15-16, as previously described^24^ and in accordance with approved procedures by the National Institutes of Health Animal Care and Usage Committee (NIDA ACUC approval# 19-OSD-10). Dams (1 animal per preparation, 8 animals total) were euthanized by isoflurane (saturated air chamber for 3-5 minutes) followed by cardiac puncture prior to removal of embryos. PCNs were plated at 6x10^4^ cells per well in polyethyleneimine-coated 96-well plates and received 50% media exchanges on day *in vitro* (DIV) 4, DIV6, DIV8, DIV11. PCNs were maintained with Neurobasal Medium (ThermoFisher Scientific, Catalog #21103049) supplemented with B27 (ThermoFisher Scientific, Catalog #17504044) and L-glutamine (Sigma-Aldrich, Catalog #8540) at 37°C with 5.5% CO_2_. Viral transductions were performed on DIV6 with the following viruses: AAV1-CaMKIIα-GLuc-ASARTDL (5.48 x 10^9^ vg/mL) or AAV1-CaMKIIα-GLuc-no tag (5.48 x 10^9^ vg/mL). On DIV8 transductions were performed for a subset of experiments with the following viruses: AAV1-CaMKIIα-KDELR1 (7.45 x 10^10^ vg/mL), AAV1-CaMKIIα-KDELR2 (7.45 x 10^10^ vg/mL), or AAV1-CaMKIIα-eYFP (7.45 x 10^10^ vg/mL). On DIV13, cells were treated with either glutamate (Sigma-Aldrich, Catalog #G8415, #G1626), thapsigargin (Tg, Sigma Aldrich, Catalog #T9033), or kainic acid (KA, Cayman Chemicals, Catalog #78050) via 50% media exchange. The vehicle for drug compounds were: 1N hydrochloric acid (Sigma-Aldrich, Catalog #320331), dimethyl sulfoxide (Sigma-Aldrich, Catalog #D8418), or phosphate buffered saline (ThermoFisher Scientific, Catalog #10010023), respectively. For a subset of experiments, cells were pre-treated with: (S)-3,5-Dihydroxyphenylglycine hydrate (DHPG; Sigma-Aldrich, Catalog #D3689), dantrolene (Sigma-Aldrich, Catalog #24868-20-0), or 2-Aminoethyl diphenylborinate (2-APB; Sigma-Aldrich, Catalog #D9754) via 20X spike-in 30 min prior to delivery of Tg or EEAs. A subset of cells was treated with glutamate or KA via 100% media exchange for time-response experiments, treatment durations are as indicated graphically. Extracellular media samples (5 µl) were collected at 24 h post-treatment. At this time, a subset of cells underwent viability testing via MTS assay (CellTiter 96® AQueous One Solution Cell Proliferation Assay, Promega, Catalog #G3582) or ATP assay (CellTiter-Glo® Luminescent Cell Viability Assay, Promega, Catalog #G7570). A subset of cells was rinsed twice with phosphate buffered saline and incubated with lysis buffer (1M Tris HCl [pH 7.5], NaCl, NP-40, and protease inhibitor cocktail [Sigma Aldrich, Catalog #P8340]) for 20 min at 4°C on a platform shaker. From this media, a 5 µl sample was collected for assay of intracellular GLuc-SERCaMP levels.

### Gaussia Luciferase (GLuc) assay

Extracellular and intracellular GLuc-SERCaMP levels were determined as previously described^19,20^. Briefly, GLuc luminescence was determined following injection of 100 µl of coelenterazine (8 µM; Regis Technologies, Catalog #1-361204-200) into white 96-well plates containing media samples, using a Synergy H1 microplate reader (Biotek; Catalog #11-120-516).

### Immunohistochemistry (IHC)

On DIV14, cells were rinsed with PBS (2X, 5 mins) and fixed with 4% paraformaldehyde. Cells were blocked in goat serum for 1h at room temperature, then incubated with mouse anti-CaMKIIα (6G9) (1:1000, Pierce Biotechnology, Catalog #MA1-048) and rabbit anti-Gaussia luciferase (1:1000, New England BioLabs, Catalog #E8023S) overnight at 4°C. Goat anti-mouse 488 (1:1000, ThermoFisher Scientific, Catalog #A-11029) and goat anti-rabbit 568 (1:1000, Catalog #A-11031) were applied for 1h at room temperature. Cells were imaged with an EVOS FL Auto 2 Imaging System (ThermoFisher Scientific, Catalog #AMAFD2000).

### Immunoprecipitation (IP)

One hundred microliters of Protein A magnetic beads (SureBeads, BioRad, Catalog #1614013) were washed with PBS + 0.1% Tween-20 (PBS-T) then incubated with 2 µL of PDI antibody (Abcam, Catalog #ab2792) in 200 µL PBS-T for 10 min (end-over-end rotation, room temperature). The beads were washed with PBS-T and 400 µL of cell culture media was incubated with the beads/antibody for 1 h (end-over-end rotation, room temperature). The beads were washed with PBS-T then eluted using 40 µL of 1x LDS (NuPage LDS, ThermoFisher Catalog #NP0007). After a 10 min incubation at 70°C, the beads were magnetized and the eluent was moved to a new tube. Twelve microliters of this eluent was loaded into a 4-12% Bis-Tris gel (ThermoFisher, 10 well: Catalog #NP0321BOX, 17 well: NP0329BOX) and run using 1x MOPS buffer (ThermoFisher Catalog #NP0001). Proteins were transferred to a 0.2 µm PVDF membrane (ThermoFisher IB24001) using an iBlot2, Program P0 (ThermoFisher Catalog # IB21001S). Blots were blocked for 1 h with Rockland blocking buffer (VWR, Catalog #RLMB-070) then incubated with PDI antibody (1:500, Abcam, Catalog #ab2792) overnight at 4°C. Goat anti-mouse IR680 (1:4000, LICOR, Catalog #925-68070) was applied for 1 h at room temperature. Blots were scanned on a LICOR Odyssey scanner (Model 9120).

### Experimental Design and Statistical Analysis

Mixed-sex primary cortical neurons were included to represent both male- and female-derived neurons in our experiments. Specific test details can be found in the results section. For experiments with two experimental groups, two-tailed t-tests were used. For experiments with three or more experimental groups, one-way ANOVA with the Dunnett’s multiple comparison *post-hoc* test was used. For experiments with two experimental conditions, two-way ANOVA with the Dunnett’s multiple comparison *post-hoc* test was used. Sample sizes were based on experience with primary cultures. All analyses were performed using GraphPad Prism 8 and values of p<0.05 were considered statistically significant. Data are represented as mean ± standard error of the mean (SEM). Any single value that was >2 standard deviations from the mean was considered outlier and omitted from the data set.

## Results

### Thapsigargin-induces GLuc-SERCaMP secretion from CaMKIIα-expressing neurons

To monitor exodosis in primary cortical neurons that respond to glutamate, we developed an adeno-associated viral vector to express our GLuc-SERCaMP reporter from the Calcium/calmodulin-dependent protein kinase II alpha promoter (AAV-CaMKIIα-GLuc-SERCaMP). We immunohistochemically confirmed expression of GLuc within CaMKIIα-positive primary cortical neurons (PCNs; **Extended Figure 1-1A).** To demonstrate functional GLuc-SERCaMP secretion in response to ER calcium depletion, PCNs were transduced with the virus then treated with thapsigargin (Tg), which specifically inhibits the SERCA pump resulting in the depletion of ER Ca^2+19,25,26^. Extracellular luciferase activity from GLuc-SERCaMP was measured at several time points following Tg and exhibited an effect of Tg (F_(4,72)_=134.2, p<.0001, **Extended Figure 1-1B**), an effect of time (F_(2,72)_=411.8, p<.0001), and an interaction between the two variables (F_(8,72)_=29.13, p<.0001). At 2 h post-treatment, Tg at 30 nM and 100 nM significantly increased extracellular GLuc-SERCaMP levels (t(72)=2.688, p=.03; t(72)=2.575, p=.04, respectively). At 4 h post-treatment, 10 nM, 30 nM, and 100 nM Tg significantly increased GLuc-SERCaMP secretion (t(72)=3.162, p=.008; t(72)=8.837, p<.0001; t(72)=9.267, p<.0001, respectively). At 24 h, 10 nM, 30 nM, and 100 nM Tg significantly increased GLuc-SERCaMP secretion (t(72)=8.606, p<.0001; t(72)=17.97, p<.0001; t(72)=18.19, p<.0001, respectively). We also measured intracellular levels of GLuc-SERCaMP 24 h after treatment with 100 nM Tg. Consistent with an increase in secretion and corresponding loss from within the cell, intracellular GLuc-SERCaMP activity were decreased following Tg treatment (t(8)=8.411, p=.0002, **Extended Figure 1-1C**). These data are consistent with previous work showing AAV vectors expressing GLuc-SERCaMP from other promoters (CMV and EF1a) are capable of reporting on exodosis^13,27,28^. CaMKIIα is expressed in glutamate-responsive neurons and actively participates in glutamate-related signaling^29^. We will use AAV-CaMKIIα-GLuc-SERCaMP to examine the effects of excitatory amino acids (EEAs) on exodosis from CaMKIIα-positive neurons.

### Excitatory amino acids trigger exodosis and reduce cell viability in a dose-dependent manner

Rat primary cortical neurons were transduced with AAV-CaMKIIα-GLuc-SERCaMP and treated with excitatory amino acid glutamate or kainic acid (KA), a glutamate analog. At 24 h post-treatments, glutamate increased secretion of GLuc-SERCaMP (F_(3,32)_=37.45, p<.0001, **Figure 1A**) in a dose-dependent manner vs. vehicle (30 µM: t(32)=3.681, p=.0024; 60 µM: t(32)=6.727, p<.0001; 100 µM: t(32)=10.15, p<.0001). Similarly, KA increased GLuc-SERCaMP secretion (F_(3,32)_=31.05, p<.0001, **Figure 1B**) in a dose-dependent manner vs. vehicle at 24 h post-treatment (30 µM: t(32)=1.794, p=.1954; 60 µM: t(32)=5.39, p<.0001; 100 µM: t(32)=8.876, p<.0001). Consistent with the ability of these EEAs to induce GLuc-SERCaMP secretion, intracellular levels of GLuc-SERCaMP were reduced following glutamate treatment (F_(3,8)_=4.919, p=.0318; **Extended Figure 1-1D**) or KA (F_(3,8)_=30.1, p=.0001, **Extended Figure 1-1F**). Post-hoc tests revealed that intracellular GLuc-SERCaMP levels were significantly reduced in response to treatment with 100 µM glutamate or KA (t(8)=3.611, p=.017; t(8)=4.546, p=.0049, respectively). Interestingly, KA increased intracellular GLuc-SERCaMP at 30 µM (t(8)=4.759, p=.0037), which suggests a potential increase in production of GLuc-SERCaMP at a dose of KA that does not promote secretion. Cell viability, as assessed by MTS assay, was reduced following treatment with glutamate (F_(3,8)_=16.69, p=.0008, **Figure 1C**), post-hoc analysis revealed a significant effect of 100 µM (t(8)=6.359, p=.0006). Kainic acid also significantly reduced viability by MTS assay (F_(3,8)_=11.08, p=.0032, **Figure 1D**), and this effect was carried by the 60 µM (t(8)=3.085, p=.037) and 100 µM doses of KA (t(8)=4.598, p=.0045). There was a positive correlation between extracellular GLuc-SERCaMP and MTS-associated viability in cell treated with glutamate (r^2^=.6163, p=.0025, **Extended Data Figure 1-1E**) or with KA (r^2^=.6944, p-value=.0008, **Extended Data Figure 1-1G**).

**Figure 1.**
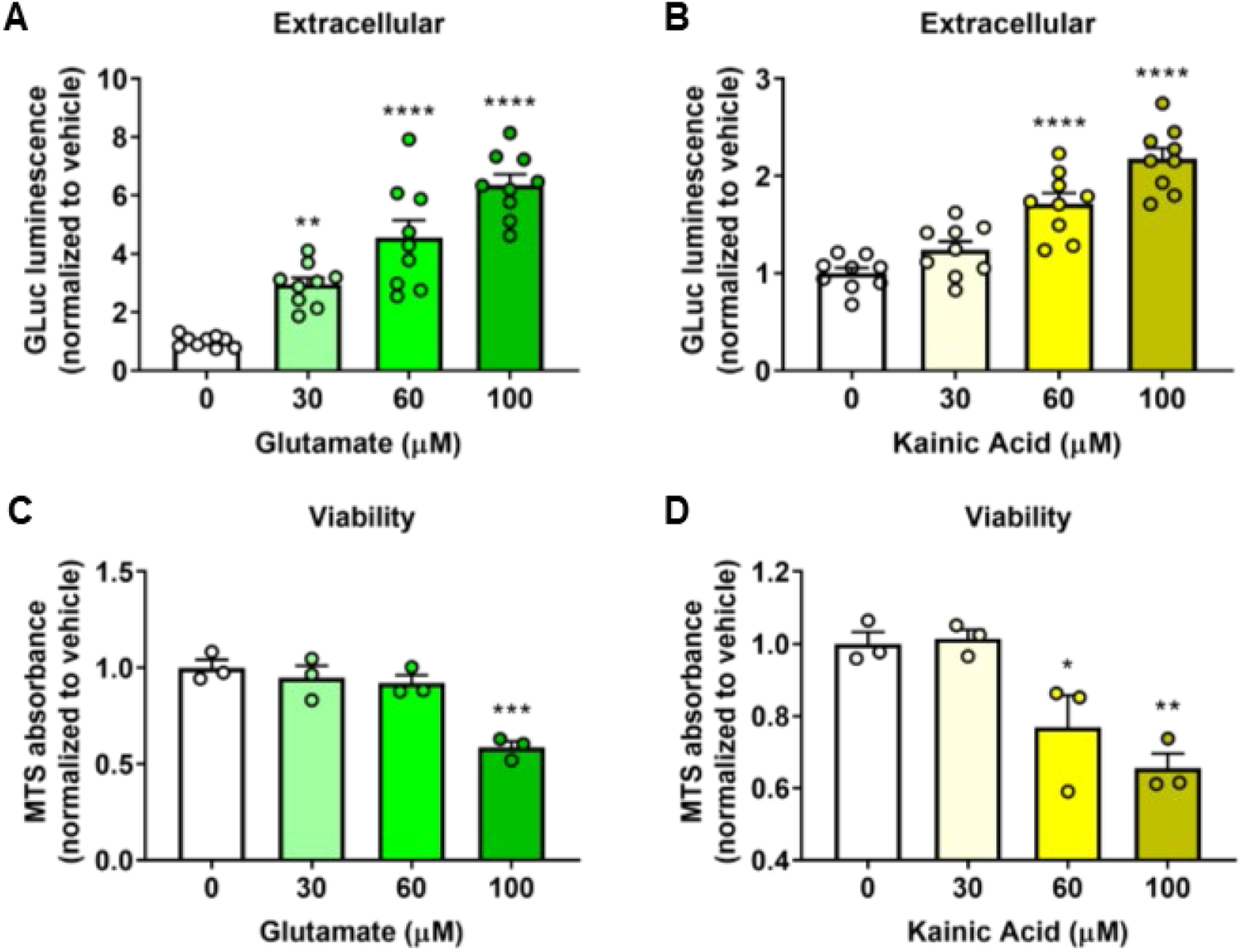
Excitatory amino acids increase extracellular GLuc-SERCaMP in a dose-dependent manner. **A)** Extracellular GLuc-SERCaMP activity following treatment with glutamate (n=9/group) and **B)** kainic acid (KA) (n=9/group). Assay of cell viability 48 h following treatment with **(C)** glutamate (n=3/group) and **(D)** KA (n=3/group), respectively. Data are shown as mean + SEM. *p<.05, **p<.01, ***p<.001, ****p<.0001 vs. vehicle.

### Pharmacological blockade of plasma membrane Ca^2+^ channels and ER Ca^2+^ channels mitigate GLuc-SERCaMP secretion induced by ER Ca^2+^ -depleting agents

Glutamate activates N-methyl-D-aspartate (NMDA) receptors to make them highly permeable to Ca^2+^ which can influence intracellular Ca^2+^ release, cell signaling and synaptic strength. To assess the contribution of NMDA receptors towards EEA-induced GLuc-SERCaMP secretion we used the NMDA receptor antagonist MK801. We first conducted a dose-response experiment to determine an effective dose of MK801 in our paradigm. Cells were incubated with MK801 at 0, 3, 10, 30, or 100 µM, 30 min prior to glutamate treatment (100 µM). MK801 was able to attenuate glutamate-induced GLuc-SERCaMP secretion (F_(4,55)_=10.84, p<.0001, **Extended Data Figure 2-2A**); with 3, 10, and 30 µM doses of MK801 blunting glutamate-induced GLuc-SERCaMP secretion as compared to glutamate alone (t(55)=4.856, p<.0001; t(55)=4.04, p=.0007; t(55)=3.384, p=.0053; respectively). Intracellular levels of GLuc-SERCaMP were not affected by MK801 in glutamate-treated cells (one-way ANOVA: F_(4,25)_=1.117, p=.37, **Extended Data Figure 2-2B**). There was a main effect of MK801 to increase viability in glutamate-treated cells (F_(4,25)_=2.897, p=.04, **Extended Data Figure 2-2C**), with an effect at the 30 µM dose compared to vehicle (t(25)=2.77, p=.0416).

After we established an effective dose of MK801, cells were incubated with 3 µM MK801 for 30 min prior to glutamate treatment (100 µM). We observed a main effect of treatment (F_(3,20)_=862.6, p<.0001, **Figure 2A**). When delivered alone, MK801 increased extracellular GLuc-SERCaMP levels as compared to vehicle (t(20)=3.708, p=.0055). Consistent with other experiments within this study, glutamate induced robust GLuc-SERCaMP secretion (t(20)=42.68, p<.0001) and MK801 blocked this effect (vs. vehicle: t(20)=.1426, p=.9998; vs. glutamate: t(20)=42.53, p<.0001), indicating that activation of excitatory NMDA receptors contributes to secretion of ER resident proteins. We found an effect of MK801 treatment on cell viability (F_(3,20)_=3.626, p=.0308, **Figure 2B**). As compared to glutamate alone, cells pretreated with MK801 prior to glutamate exhibited higher viability (t(20)=3.05, p=.0251). We observed similar effects of NDMA receptor blockade on KA-induced ER Ca^2+^ depletion. There was a main effect of MK801 to mitigate KA-induced (100 µM) GLuc-SERCaMP secretion (F_(4,55)_=11.93, p<.0001, **Extended Data Figure 2-2D**); where 3 and 10 µM MK801 lowered extracellular GLuc-SERCaMP vs. vehicle (t(55)=4.088, p=.0006; t(55)=2.808, p=.0276, respectively). MK801 reduced intracellular GLuc-SERCaMP levels (F_(4,25)_=5.937, p=.0017, **Extended Data Figure 2-2E**), carried by 100 µM KA vs. vehicle (t(25)=2.783, p=.0405). MK801 also increased viability in KA-treated cells (F_(4,25)_=3.379, p=.024, **Extended Data Figure 2-2F**), with an effect at 10 and 30 µM MK801 vs. vehicle (t(25)=2.882, p=.032; t(25)=3.147, p=.0169; respectively). Collectively, these data support that EEA-induced Ca^2+^ influx leads to the secretion of ER resident proteins

**Figure 2.**
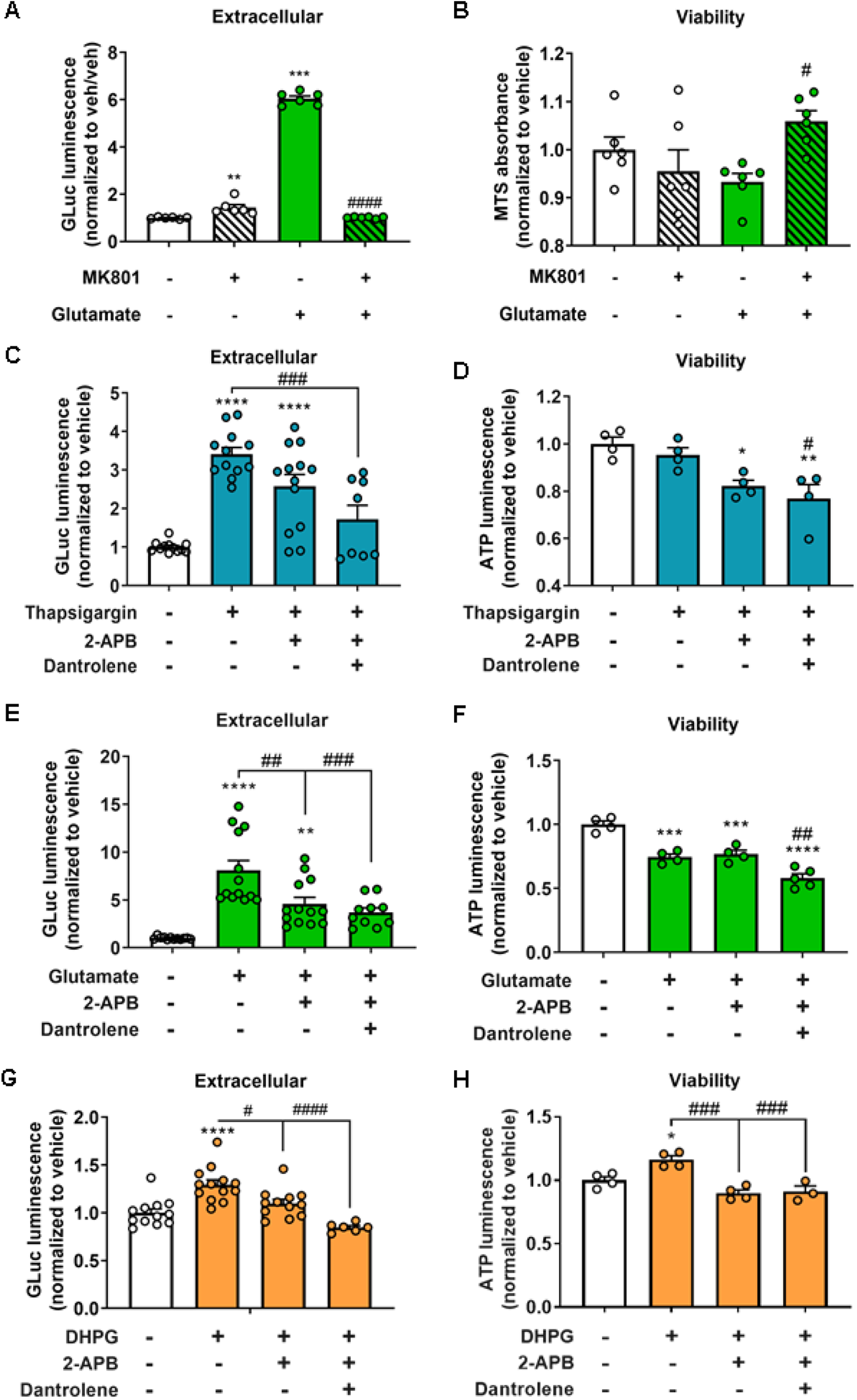
Pharmacological blockade of NMDA receptor and ER Ca^2+^ efflux channels attenuate GLuc-SERCaMP secretion induced by Tg and glutamate. **A)** MK801 (3 µM) added to 30 min prior to 24 hr glutamate (100 µM) exposure blocks GLuc-SERCaMP secretion (n=6/group) and **(B)** cell viability (n=6/group). **C)** Effect of 2-APB with or without dantrolene on Thapsigargin (Tg, 100 nM)-induced GLuc-SERCaMP secretion 24 h post-treatment (n=8-13/group) and **(D)** cell viability (n=4/group) **E)** Effect of 2-APB (IP_3_R antagonist, 50 µM) with or without Dantrolene (RyR antagonist, 30 µM) on glutamate-induced GLuc-SERCaMP secretion (n=10-13/group) and **(F)** cell viability (n=6/group). **G)** Effect of DHPG (50 nM) on GLuc-SERCaMP secretion 24 h post-treatment (n=6-13/group) and **(H)** viability (n=3-4/group). Data are shown as mean + SEM. **p<.01, ****p<.0001 vs. vehicle; #p<.05, ##p<.01, ###p<.005, ####p<.0001 vs. glutamate, Tg, or DHPG alone.

Within the cell, 1,4,5-triphosphate receptors (IP_3_R) and ryanodine receptors (RyR) control the efflux of Ca^2+^ from the ER^30^. To examine the role of these ER Ca^2+^ channels in ER Ca^2+^ depletion, we first tested the effects of modifying ER channels on exodosis caused by Tg. There was a main effect of Tg (100 nM; F_(3,41)_=20.56, p<.0001, **Figure 2C**), and when delivered alone, Tg increased extracellular GLuc-SERCaMP (t(41)=7.468, p<.0001). After confirming the functionality of this pharmacological tool, we confirmed the ability of these doses of 2-APB and dantrolene to mitigate effects of Tg to induce GLuc-SERCaMP secretion, in which we pre-treated cells with an IP_3_R antagonist (2-APB, 50 µM), with or without co-incubation with a RyR antagonist (dantrolene, 30 µM). As compared to vehicle, incubation with 2-APB did not completely block the effects of Tg (t(41)=4.994, p<.0001), yet there was a trend toward a reduction in extracellular GLuc-SERCaMP as compared to Tg alone (t(41)=2.662, p=.061). The 2-APB plus dantrolene cocktail reduced Tg-induced GLuc-SERCaMP secretion as compared to Tg alone (t(41)=4.682, p=.0002). There was a main effect of treatment condition on cellular viability (F_(3,12)_=8.195, p=.0031, **Figure 2D**). Post-hoc analysis revealed that as compared to vehicle, Tg alone did not change viability (t(12)=.8682, p>.9999). However, cells treated with Tg plus 2-APB (t(12)=3.315, p=.0308) and Tg plus 2-APB and dantrolene (t(12)=4.311, p=.0051) exhibited reduced viability as compared to vehicle-treated cells. Somewhat surprisingly, as compared to Tg alone, Tg plus 2-APB and dantrolene reduced viability (t(12)=3.443, p=.0243). These data support that IP_3_R and RyR contribute to Tg-induced exodosis in CaMKIIa neurons and is consistent with our previous studies^13,27,31^. However, at the doses employed, the antagonists for these ER-associated receptors were not sufficient to protect against Tg-induced decrements in cellular viability. The role of IP3R and RyR in EEA-induced exodosis were probed using cotreatment of glutamate with 2-APB (50 µM) along with or without dantrolene (30 µM). There was a main effect of glutamate (100 µM) to increase extracellular GLuc-SERCaMP levels (F_(3,44)_=18.67, p<.0001, **Figure 2E**), and when delivered alone, glutamate increased extracellular GLuc-SERCaMP vs. vehicle (t(44)=7.397, p<.0001). Co-treatment with 2-APB and glutamate reduced GLuc-SERCaMP secretion as compared to glutamate alone (t(44)=3.724, p=.0028). The combination treatment of 2-APB and dantrolene also blunted glutamate-induced GLuc-SERCaMP secretion (t(44)=4.369, p=.0004), but did not show additive effect compared to 2-APB alone despite approaching vehicle treated levels (t(44)=2.624, p=.0595). Examination of cellular viability revealed a main effect of glutamate treatment (F_(3,13)_=34.24, p<.0001, **Figure 2F**). There was no protection against glutamate-induced reductions in viability when compared to vehicle (glutamate: t(13)=5.84, p=.0003; glutamate + 2-APBL t(13)=5.317, p=.0007; glutamate + 2-APB and dantrolene: t(13)=10.12, p<.0001). Further, cells treated with the combination of 2-APB, dantrolene, and glutamate exhibited reduced viability vs. cells treated with glutamate alone (t(13)=3.96, p=.0082). These data show that IP3R and RyR activation contribute to glutamate-induced exodosis, yet antagonism of these ER-associated receptors is not sufficient to protect cells against glutamate-induced reductions in viability.

Metabotropic glutamate receptors such as mGluR1 can promote release of ER-calcium via activation of IP_3_R^32^ and we show above that IP3R antagonism can attenuate glutamate-mediated exodosis. We next tested whether metabotropic glutamate receptor activation alone was capable of triggering exodosis. Treatment of primary neurons expressing GLuc-SERCaMP with 3,5 dihydroxyphenylglycine (DHPG, 50 µM), a group 1 metabotropic glutamate receptor agonist, resulted in a main effect of DHPG treatment (F_(3,39)_=15.0, p<.0001, **Extended Data Figure 2-G**). DHPG significantly increased extracellular GLuc-SERCaMP levels vs. vehicle (t(39)=4.942, p<.0001). As compared to DHPG alone, cells pretreated with 2-APB and 2-APB plus dantrolene blunted GLuc-SERCaMP secretion (t(39)=3.301, p=.0103; t(39)=6.103, p<.0001, respectively). The drugs without agonist had no effect. Viability was also affected by treatment (F_(3,11)_=15.67, p=.0003, **Figure 2H**). Post-hoc analysis revealed that DHPG increased viability vs. vehicle (t(12)=3.822, p=.0142). The data indicate that metabotropic glutamate receptor activation can cause exodosis but to a lesser extent compared to ionotropic glutamate receptor activation (see Figure 2A above).

### KDELR1 and KDELR2 diminish Tg-induced GLuc-SERCaMP secretion

ERS tails of proteins that move from ER to Golgi interact with KDEL receptors located in the Golgi and cause retrograde transport of the ERS-containing proteins from the Golgi back to the ER. We have previously shown that augmenting KDELR expression can attenuate exodosis caused by thapsigargin, hypoxia and hyperthermia^13,33^. We first investigated the ability of KDELR to influence Tg-induced GLuc-SERCaMP secretion in rat primary neurons transduced with AAV vectors expressing GLuc-SERCaMP and eYFP (control), KDELR1, or KDELR2. There was a main effect of Tg (100 nM; F_(2,135)_=36.58, p<.0001, **Figure 3A**) and of KDELR overexpression (F_(2,135)_=20.21, p<.0001). Post-hoc examination revealed Tg induced GLuc-SERCaMP secretion only in control cells (control: t(135)=8.648, p<.0001; KDELR1: t(135)=.03458, p>.9999; KDELR2: t(135)=1.89, p=.172). Additionally, KDELR1-and KDELR2 overexpression mitigated Tg-induced GLuc-SERCaMP secretion as compared to control (t(135)=8.612, p<.0001; t(135)=6.665, p<.0001; respectively). Comparisons revealed no difference between the KDELR isoforms (t(135)=1.855, p=.9982). Examination of viability revealed no effect of transduction condition on viability (F_(2,52)_=.523, p=.5958, **Figure 3C**), but there was a trend for Tg to reduce viability (F_(1,52)_=3.655, p=.0614). These data show that overexpression of both isoforms of the KDELR prevent release of GLuc-SERCaMP suggesting they attenuate exodosis caused by ER calcium depletion in neurons. Through these experiments, we established a paradigm for determining the effectiveness of these KDELR isoforms to prevent glutamate-induced exodosis.

**Figure 3.**
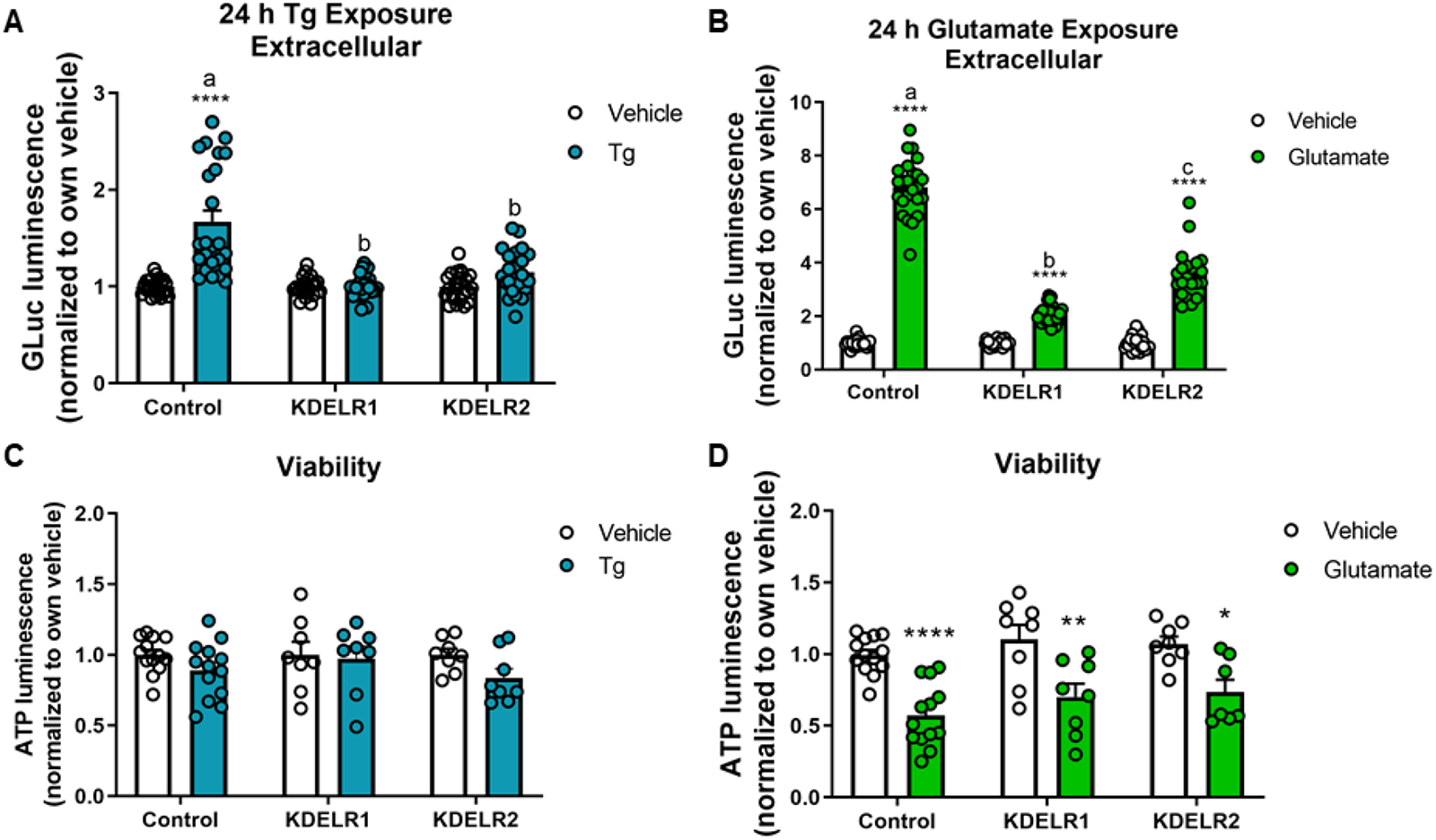
Augmenting KDELR expression mitigates GLuc-SERCaMP secretion in KDELR isoform-dependent manner. **A)** AAV-mediated delivery of KDELR1 and KDELR2 to rat primary neurons reduces thapsigargin (Tg, 100 nM)-induced GLuc-SERCaMP secretion (n=31-42/group) and (**B)** does not affect cell viability (n=8-13/group). **C)** Increased expression of KDELR1 and KDELR2 decreases glutamate (100 µM)-induced GLuc-SERCaMP secretion 24 h post-treatment (n=34-42/group). **D)** KDELR expression does not affect viability (n=7-13/group). Data are shown as mean + SEM. **p<.01, ****p<.0001 vs. vehicle. Different letters indicate statistically significant differences between groups.

### KDELR1 overexpression dampens glutamate- and DHPG-induced SERCaMP secretion

After confirming the ability of two KDELR isoforms to mitigate Tg-induced GLuc-SERCaMP secretion, we assessed their impact on glutamate-induced secretion. We observed a main effect of treatment (100 µM, F_(1,135)_=1021.0, p<.0001, **Figure 3B**) and KDELR condition (F_(2,135)_=202.3, p<.0001). Glutamate increased extracellular GLuc-SERCaMP in all conditions (control: t(135)=34.32, p<.0001; KDELR1: t(135)=6.483, p<.0001; and KDELR2: t(135)=14.68, p<.0001). Comparisons across KDELR conditions revealed that KDELR1-and KDELR2-overexpression blunted secretion vs. control (t(135)=27.83, p<.0001; t(135)=18.48, p<.0001; respectively). In glutamate treated cells, KDELR1 more effectively blunted GLuc-SERCaMP secretion vs. KDELR2 (t(135)=8.414, p<.0001). Post hoc analysis revealed an effect of glutamate to induce GLuc-SERCaMP secretion in cells expressing control, KDELR1, and KDELR2 (t(51)=5.874, p<.0001; t(51)=3.167, p=.0078; t(51)=2.725, p=.0261; respectively).There was a main effect of glutamate to reduce viability (F_(1,51)_= 41.78, p<.0001, **Figure 3D**), yet there was no effect of KDELR overexpression (F_(2,51)_= 1.171, p=.3183). Collectively, we show that KDELRs are capable of attenuating glutamate-induced exodosis.

### Impact of EEAs on a constitutively secreted GLuc and cell viability

To rule out the observed effects with GLuc-SERCaMP were not due to overall changes in secretion, we used “GLuc-no tag” which does not contain a carboxy-terminal ERS and thus is not retained in the ER but rather is constitutively secreted from the cell in a manner independent of ER Ca^2+^ depletion^19^. Glutamate significantly decreased the extracellular activity of this control reporter (F_(3,32)_=32.48, p<.0001, **Figure 4A**) vs. vehicle (30 µM: t(32)=5.593, p<.0001; 60 µM: t(32)=7.882, p<.0001; 100 µM: t(32)=9.008, p<.0001). Similarly, KA reduced extracellular levels of GLuc-no tag (F_(3,32)_=64.64, p<.0001; **Figure 4B**) vs. vehicle (30 µM: t(32)=2.591, p=.0377; 60 µM: t(32)=7.723, p<.0001; 100 µM: t(32)=12.82, p<.0001). There was no effect of glutamate to reduce intracellular levels of GLuc-no tag (**Figure 4C**). However, there was a main effect of KA to reduce intracellular GLuc (F_(3,8)_=215.5, p<.0001; **Figure 4D**), which was bidirectional in nature. The finding that EEAs reduce secretion of GLuc-no tag are in direct contrast to what was observed with GLuc-SERCaMP (enhanced secretion). Taken together, these data support that EEAs actively induce secretion of ERS proteins rather than promote cause general increase in protein secretion.

**Figure 4.**
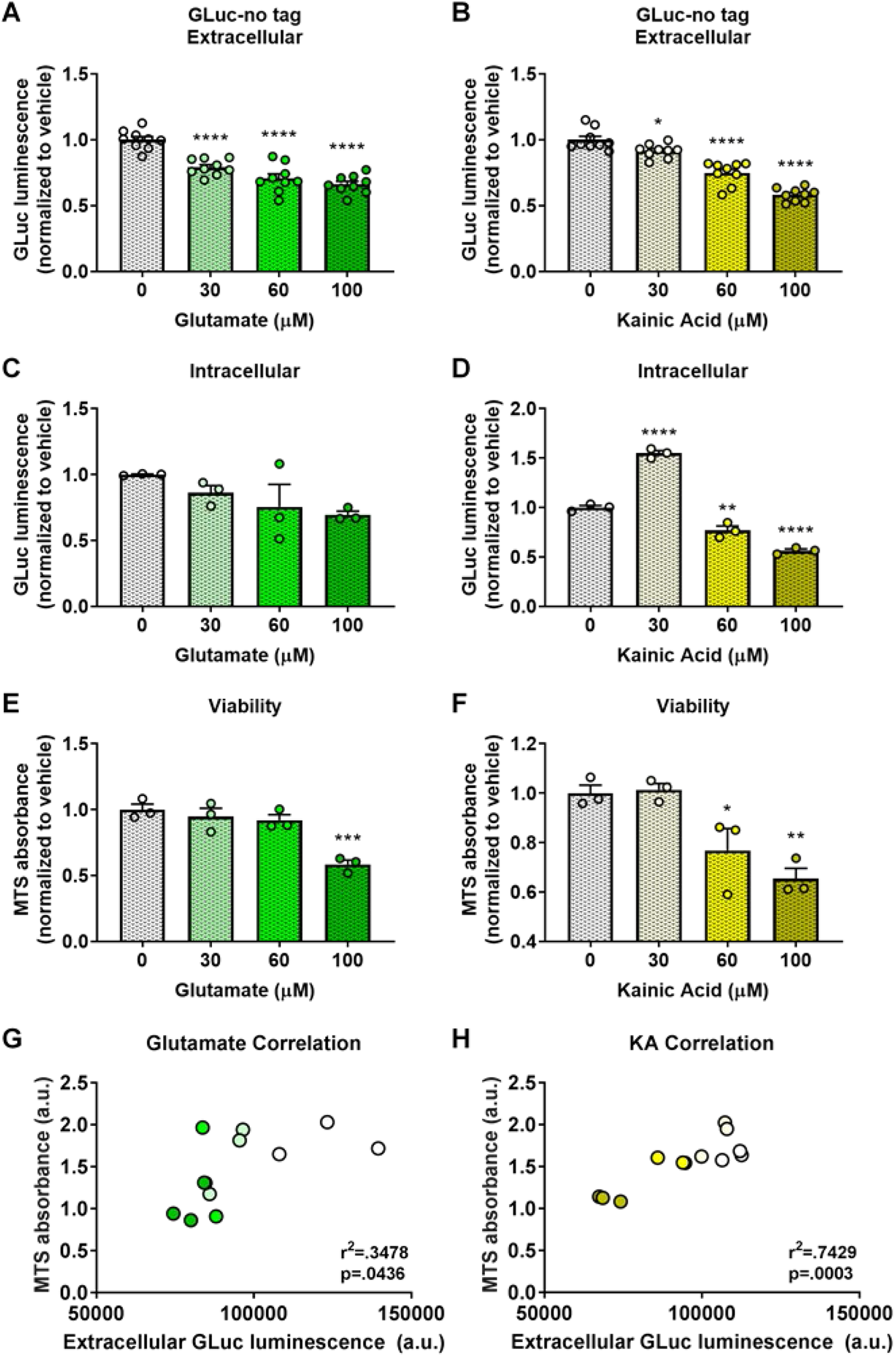
Effects of EEAs on the constitutively secreted GLuc-no tag and cell viability. **A)** Glutamate (30, 60, and 100 µM) and (**B)** kainic acid (KA, 60 and 100 µM) reduced secretion of GLuc-no tag 24 h post-treatment (n=9/group). **C)** There was no significant effect of glutamate on intracellular GLuc-no tag levels. **D)** In contrast, KA at 30 µM increased and 60 and 100 µM significantly reduced intracellular levels of GLuc-no tag. **E)** Glutamate (100 µM) and (**F)** KA (60 and 100 µM) significantly reduced viability 48 h post-treatment, and there was a significant correlation between (**G)** glutamate-and (**H)** KA-induced secretion of GLuc-SERCaMP and viability (n = 3/group, Pearson correlation). Data are shown as mean + SEM. n = 9/group; *p<.05, **p<.005, ***p<.001, ****p<.0001 vs. vehicle.

There was a main effect of glutamate to reduce viability of cells expressing GLuc-no tag (MTS assay: F_(3,8)_=16.69, p=.0008, **Figure 4E**), with the 100 µM concentration carrying the effect (t(8)=6.359, p=.0006). KA also reduced viability (F_(3,8)_=11.08, p=.0032; **Figure 4F**), with 60 µM and 100 µM carrying the effect (t(8)=3.085, p=.0369; t(8)=4.598, p=.0045, respectively). There was a significant correlation between secreted levels of GLuc-no tag and MTS-associated viability in glutamate-treated cells (r^2^=.6163, p=.0025; Figure 4G) and in KA-treated cells (r^2^=.6944, p-value=.0008, **Figure 4H**). These correlations reveal that lower viability is associated with reduced GLuc-no tag secretion; which contrasts with the correlation observed for GLuc-SERCaMP in which lower viability was associated with increased secretion. Together these data indicate that compromised cell viability associated with glutamate treatment does not trigger general protein secretion.

### Time-dependent effect of EEAs to increase GLuc-SERCaMP secretion and decrease GLuc-no tag secretion

As physiological levels of glutamate are elevated for relatively brief periods of time in disease states such as stroke^34^ and seizure, we assessed the effect of acute glutamate exposure on exodosis related to ER Ca^2+^ depletion. There was a significant effect of glutamate exposure (100 µM) to increase secretion of GLuc-SERCaMP (F_(8,164)_=34.66, p<.0001, **Figure 5A**), with effects observed with as little as 2 min exposure (120 sec: t(164)=3.628, p=.0028; 300 sec: t(164)=6.086, p<.0001; 600 sec: t(164)=5.906, p<.0001; 1800 sec: t(164)=10.33, p<.0001; and 24 h: t(164)=12.62, p<.0001). Consistent with increased secretion of GLuc-SERCaMP, intracellular GLuc-SERCaMP was reduced in a time-responsive fashion (F_(8,72)_=7.944, p<.0001, **Figure 5B**). Reductions in intracellular GLuc-SERCaMP were observed with as little as 1 min of glutamate exposure (60 sec: t(72)=3.502, p=.0059; 120 sec: t(72)=4.175, p=.0007; 300 sec: t(72)=4.341, p=.0004; 600 sec: t(72)=3.649, p=.0037; 1800 sec: t(72)=5.238, p<.0001; and 24 h: t(72)=6.695, p<.0001). We also observed a reduction in viability following glutamate treatment (F_(8,54)_=25.69, p<.0001, **Figure 5C**), with significant effects emerging from 5 min of exposure (300 sec: t(54)=5.724, p<.0001; 600 sec: t(54)=5.489, p<.0001; 1800 sec: t(54)=8.465, p<.0001; and 24 h: t(54)=8.574, p<.0001). These data indicate that in addition to promoting ER Ca^2+^ depletion following protracted exposure (e.g., 24 h), a physiologically relevant timeline of glutamate exposure is sufficient to induce exodosis.

**Figure 5.**
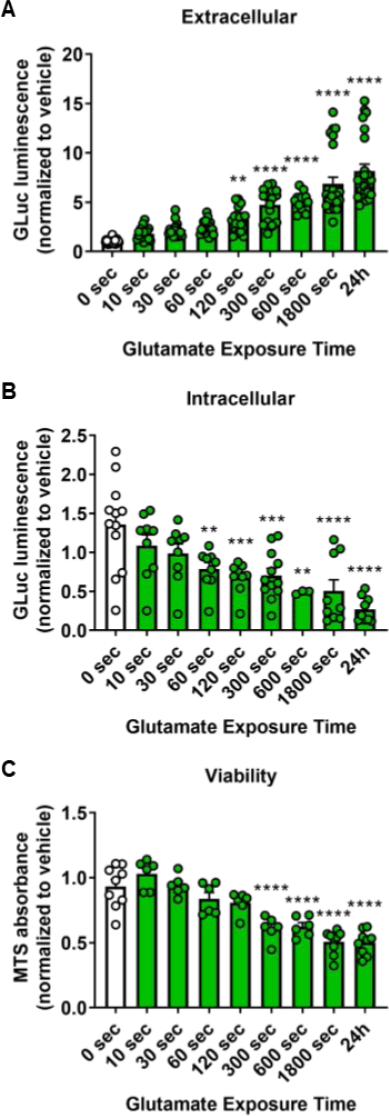
Duration of EEA exposure impacts degree of GLuc-SERCaMP secretion and viability in a time-dependent manner. **A)** Extracellular GLuc luminescence following treatment with glutamate (100 µM) for ascending durations (n=12-24/group). **B)** Intracellular GLuc luminescence following treatment with glutamate (n=3-12/group). **C)** Assay of cellular viability 24 h following exposure to glutamate (100 µM) (n=6-9/group). Data are normalized to time-matched vehicle-treated cells, shown as mean + SEM. *p<.05, **p<.005, ***p<.001, ****p<.0001 vs. 0 min.

Consistent with our glutamate data, KA (100 µM) also induced GLuc-SERCaMP secretion in a time-dependent fashion (F_(7,40)_=274.4, p<.0001, **Extended Data Figure 5-5A**). KA increased GLuc-SERCaMP secretion with as little as 5 min of exposure (t(40)=6.276, p<.0001) and with 30 min and 24 h exposure (t(40)=26.59, p<.0001; t(40)=28.09, p<.0001, respectively). The duration of KA exposure influenced viability (F_(7,16)_=84.94, p<.0001). Post-hoc analysis revealed a significant reduction in viability measured by total ATP following 1 min, 5 min, and 24 h exposure (1 min: t(16)=3.765, p=.009; 300 sec: t(16)=4.745, p<.0001; 1800 sec: t(16)=15.56, p<.0001; and 24 h: t(16)=15.22, p<.0001; **Extended Data Figure 5-5B**).

To control for effects of acute EEAs on general secretion, we employed GLuc-no tag as our control. There was a main effect of glutamate (100 µM) to reduce secretion of GLuc-no tag (F_(8,117)_=79.22, p<.0001, **Extended Data Figure 5-5C**), with significant effects observed as early as 10 sec of exposure and every time point examined thereafter (10 sec: t(117)=3.161, p=.0143; 30 sec: t(117)=5.152, p<.0001; 1 min: t(117)=6.941, p<.0001; 2 min: t(117)=10.29, p<.0001; 5 min: t(117)=13.28, p<.0001; 10 min: t(117)=14.75, p<.0001; 30 min: t(117)=18.43, p<.0001; and 24 h: t(117)=17.93, p<.0001). This effect was not due to enhanced retention of the GLuc-no tag reporter, as intracellular GLuc-no tag showed a decrease, not increase, by glutamate (100 µM; (F_(8,51)_=3.73, p=.0017, **Extended Data Figure 5-5E**). Significant effects were observed as early as 1 min of exposure and every time point examined thereafter (1 min: t(51)=2.896, p=.0373; 2 min: t(51)=3.686, p=.0041; 5 min: t(51)=3.736, p=.0035; 10 min: t(51)=3.805, p=.0028; 30 min: t(51)=4.287, p=.0006; and 24 h: t(51)=3.358, p=.0107). In line with our observations with GLuc-SERCaMP-expressing cells, glutamate (100 µM) reduced viability in GLuc-no tag-expressing cells (F_(8,54)_=24.7, p<.0001; **Extended Data Figure 5-5D**).

### Acute, transient exposure to EEAs is sufficient to cause GLuc-SERCaMP secretion from neurons

Having established that transient glutamate treatments can cause exodosis, we assessed the contribution of NMDA receptors towards EEA-induced GLuc-SERCaMP secretion after only 30 min of glutamate exposure followed by washout for 24 hrs. We pretreated cells with MK801 (3 µM, 30 min) prior to acute treatment with glutamate (100 µM, 30 min duration) and observed a main effect of treatment (F_(3,139)_=363.9, p<.0001, **Figure 6A**). As in the previous experiment, glutamate induced GLuc-SERCaMP secretion with only 30 min of exposure (t(139)=26.12, p<.0001). When delivered alone, MK801 had no effect on extracellular GLuc-SERCaMP levels as compared to vehicle (t(139)=1.59, p=.3838). We show that MK801 blocks the effect of acute glutamate exposure to induce GLuc-SERCaMP secretion (vs. vehicle: t(139)=0.9949, p=.7881; vs. glutamate: t(139)=27.11, p<.0001) indicating that activation of NMDA receptors contributes to the secretion of ERS-containing proteins. Cell viability was also assessed (**Figure 6B**). Consistent with previous experiments within this study, glutamate reduced viability as compared to vehicle-treated cells (t(44)=4.445, p=.0002) and MK801 blocks the effect of glutamate (vs. vehicle: t(44)=1.426, p=.5044; vs. glutamate: t(44)5.871, p<.0001).

**Figure 6.**
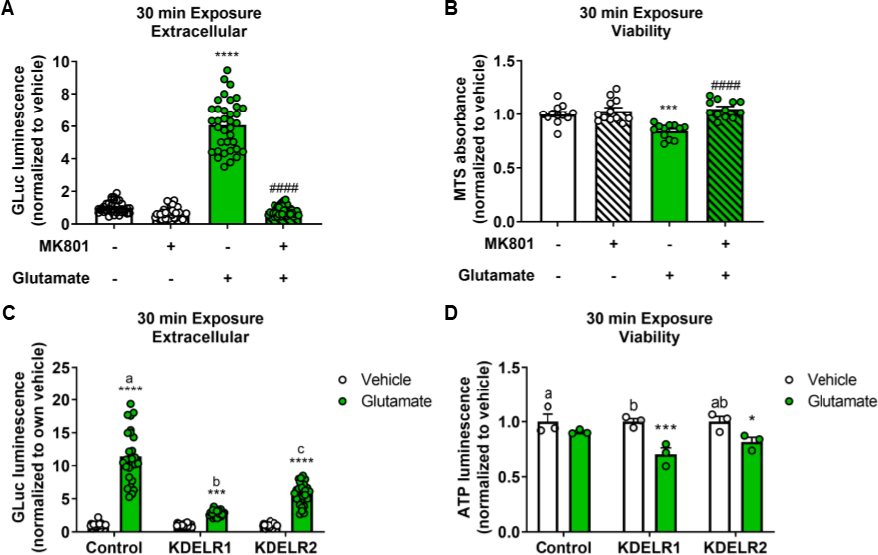
Acute glutamate exposure modulates GLuc-SERCaMP secretion. **A)** MK801 (3 µM) effects on glutamate (100 µM)-induced secretion (n=35-36/group) and **B)** viability (n=12/group). **C)** Effects of KDELR1 and KDELR2 overexpression on glutamate-induced GLuc-SERCaMP secretion (n=27/group) and **(D)** viability (n=3/group). Data are shown as mean + SEM. *p<.05, **p<.01, ***p<.001, ****p<.0001 vs. vehicle. ####p<.0001 vs. glutamate. Different letters indicate statistically significant differences between groups.

We next tested the ability of KDELR overexpression to influence GLuc-SERCaMP secretion induced by acute, transient exposure to glutamate. We observed an effect of KDELR overexpression condition (F_(2,156)_=81.23, p<.0001, **Figure 6B**) and an effect of 30 min glutamate treatment (F_(2,156)_=413.9, p<.0001) on extracellular GLuc-SERCaMP. Glutamate increased secretion in all conditions (eYFP control: t(156)=21.55, p<.0001; KDELR1: t(156)=3.853, p=.0005; KDELR2: t(156)=9.838, p<.0001). As compared to control, KDELR1 and KDELR2 overexpression blunted glutamate-induced GLuc-SERCaMP secretion (t(156)=17.7, p<.0001; t(156)=11.72, p<.0001, respectively). Further, KDELR1 overexpression more effectively blunted glutamate-induced GLuc-SERCaMP secretion as compared to KDELR2 (t(156)=5.985, p<.0001). Assessment of viability by ATP measurement in a subset of cells confirmed that glutamate reduced ATP levels (F_(1,12)_=27.37, p=.0002, **Figure 6D**) and revealed an effect of KDELR to reduce ATP levels (F_(2,12)_=4.236, p=.0406). Glutamate reduced viability in cells overexpressing KDELR1 or KDELR2 (t(12)=4.987, p=.0009; t(12)=2.858, p=.0426; respectively). In vehicle-treated cells, KDELR1 overexpression increased viability as compared to control (t(12)=2.518, p=.0084).

### Glutamate treatment promotes secretion of the endogenous ER resident protein, protein disulfide isomerase (PDI)

To extend our findings beyond the exogenous reporter GLuc-SERCaMP expressed in CaMKIIα neurons, we examined endogenous protein disulfide isomerase (PDI) secretion from all cells in our primary cortical cultures which contain non-CaMKIIα neurons and astrocytes. PDI is an endogenous ER resident protein with a C-terminal ERS tail and undergoes exodosis in response to ER calcium depletion^13^. Consistent with the GLuc-SERCaMP findings described above, western blots confirmed that glutamate (100 µM) caused the secretion of an endogenous ERS protein, PDI, in a time-dependent fashion (vs. time-matched vehicle, 0 sec: .75; 10 sec: 1.0; 30 sec: .96; 1 min: 1.41; 2 min: 1.24; 5 min: 1.32; 30 min: 2.61; 24 h: 4.01, **Figure 7A**). To assess the contribution of NMDA receptors towards EEA-induced PDI secretion, we incubated cells with MK801 (3 µM) 30 min prior to glutamate treatment (100 µM, 24 hr duration). We observed a significant main effect of treatment (F_(3,12)_=7.578, p=.0042, **Figure 7B**). MK801 alone had no effect on extracellular PDI levels as compared to vehicle (t(12)=.4393, p=.9879). Consistent with our findings using the GLuc-SERCaMP reporter, treatment with glutamate resulted in increased levels of PDI (t(12)=4.281, p=.0043). Importantly, we demonstrate that MK801 blocks the effect of glutamate to increase extracellular PDI (vs. vehicle: t(12)=.9694, p=.8231; vs. glutamate: t(12)=.5301, p=.9758). Taken together these findings indicate that activation of excitatory NMDA receptors results in the secretion of ER resident proteins.

**Figure 7.**
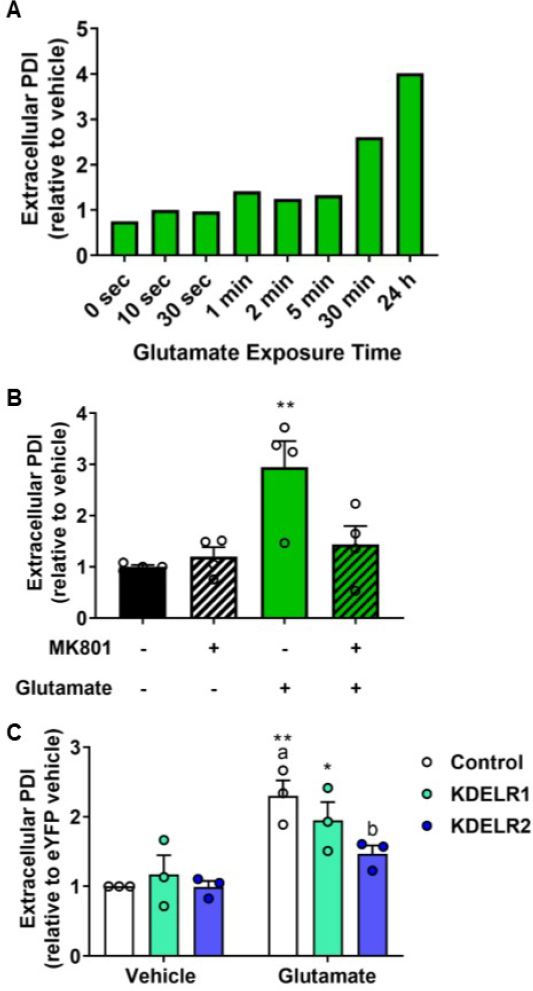
Western blot analysis of glutamate-induced secretion of the endogenous ER resident protein, phosphodiesterase (PDI). **A)** PDI secretion in response to increasing durations of glutamate exposure (n=1/group). **B)** Glutamate-induced PDI secretion and effects of MK801 pretreatment (n=4/group). **C)** Effect of KDELR overexpression on glutamate-induced PDI secretion (n=3/group). Data are shown as mean + SEM. *p<.05, **p<.005 vs. vehicle. Different letters indicate statistically significant differences between KDELR conditions.

To extend our findings of KDELRs blunting the GLuc-SERCaMP secretion in glutamate-treatment conditions, we examined the ability of KDELR to mitigate secretion of PDI. We observed an effect of glutamate treatment (F_(1,12)_=30.0, p=.0001, **Figure 7C**), yet not for KDELR condition (F_(2,12)_=2.69, p=.1083). Glutamate increased extracellular PDI in control (t(12)=4.83, p=.0012) and KDELR1 over-expression conditions (t(12)=2.893, p=.0399), but not in cells overexpressing KDELR2 (t(12)=1.763, p=.2791). In glutamate-treated cells, those in the control condition exhibited higher levels of PDI secretion as compared to KDELR2-expressing cells (t(12)=3.09, p=.0278) indicating that KDELR2 overexpression could reduce the loss of an endogenous ERS protein.

## Discussion

Dysregulation of synaptic glutamate levels can lead to excitotoxicity as is observed in conditions like stroke, traumatic brain injury, and epilepsy. The role of Ca^2+^ at the plasma membrane and within the cytoplasm towards the development of excitotoxicity is well established. However, less is known regarding the impact of glutamate mishandling on ER-Ca^2+^-mediated processes such as proteostasis. The present study demonstrates that EEAs influence ER proteostasis, specifically in CaMKII-expressing excitatory neurons. Using a pharmacological model of excitotoxicity, we found evidence of GLuc-SERCaMP secretion during excitotoxicity. In previous work, we show that ER Ca^2+^ depletion caused by thapsigargin leads to GLuc-SERCaMP secretion^19,25^. Here, we present data that a physiologically relevant regulator of intracellular depletion, i.e. glutamate, causes the secretion of ER resident proteins which can be blocked by stabilizing ER calcium. Specifically, we show that glutamate triggers the exodosis phenomenon^13^. We found that EEA-induced GLuc-SERCaMP secretion was blunted by an NMDAR antagonist, supporting that EEA-induced Ca^2+^ influx contributes to changes in ER Ca^2+^ homeostasis and proteostasis. Pharmacological stabilization of ER Ca^2+^ attenuated the EEA-induced secretion of our GLuc-SERCaMP reporter, providing evidence that ER Ca^2+^ depletion is required for EEA-induced exodosis. Further, we demonstrate that overexpressing KDELR1 blunted the EEA-induced shift in the ER proteome, supporting the hypothesis that EEA-induced exodosis results from an impaired or overwhelmed KDELR retrieval pathway. Lastly, we show that an endogenous ERS protein, PDI, undergoes secretion in response to EEA which further supports that excessive EEAs trigger ER calcium decreases and loss of ER resident proteins (i.e. exodosis).

The EEAs, glutamate and KA promote hypersynchronous neuronal activity and excitotoxicity^35–38^ and rapidly increase levels of cytosolic Ca^2+7,39–41.^ The increased Ca^2+^ plays a critical role in the cell death associated with glutamate-induced excitotoxicity^42^. We demonstrated that EEAs increased extracellular levels of GLuc-SERCaMP, which is consistent with a depletion of ER Ca^2+^ and disruption of the ER proteome^13,19^. It is notable that 2 min of glutamate exposure was sufficient to induce GLuc-SERCaMP secretion, congruent with the temporal profile of synaptic glutamate that is observed during a seizure^43^. The opposite effect on extracellular GLuc activity was observed with acute exposure to glutamate using the GLuc-no tag construct, which lacks an ERS and is constitutively secreted. The decreased secretion of the GLuc-no tag construct in response to glutamate indicates that general protein secretion or transcription of reporter does not account for our observed increases in GLuc-SERCaMP^19,31^. Taken together, our data support the connection between EEAs, ER Ca^2+^ release, and secretion of ER resident proteins as a component of excitotoxicity.

IP_3_R and RyR are primary regulators of Ca^2+^ efflux from the ER lumen to the cytosol. Activation of mGluR1,5 generates IP_3_ activity at the IP_3_R^32^ and induces ER Ca^2+^ depletion, while antagonism of IP_3_R (e.g., 2-APB) effectively blunts ER Ca^2+^ depletion^19,44^. RyR-mediated Ca^2+^ efflux is achieved via multiple effectors, including Ca^2+^ and calmodulin^45^. Dantrolene antagonizes RyR and is an effective treatment for conditions such as malignant hyperthermia^46^. In the present study, Tg and EEA-induced GLuc-SERCaMP secretion was blunted by antagonism of IP_3_R, and further decreased when given in combination with a RyR antagonist. Dantrolene preferentially acts at RyR1 and RyR3 isoforms^47^, while RyR2 is the predominant isoform in neurons^48^. This is consistent with the present study in which we observed little to no effects of dantrolene when administered alone, prior to EEAs. Our results further support that EEAs promote ER Ca^2+^ efflux.

NMDAR plays a key role in synaptic communication and plasticity primarily through their ability to allow extracellular Ca^2+^ influx^49^, which can also contribute to the development of excitotoxicity. We examined the ability of an NMDAR antagonist to mitigate an EEA-induced shift in the ER proteome. Pre-treatment with MK801, an NMDAR antagonist, significantly blunted glutamate-induced secretion of GLuc-SERCaMP. MK801 also protected cells against glutamate-induced changes in viability however the contribution of exodosis to excitotoxicity requires further investigation. Taken together, these data support that EEAs induce GLuc-SERCaMP secretion via increased activity of the NMDAR to change intracellular calcium. Moreover, our results support that excitotoxic conditions decrease ER Ca^2+^ with a concurrent secretion of ER resident proteins i.e. exodosis. Further studies are needed to better understand how the redistribution of the ER proteome contributes to changes in neuronal function and the surrounding cells exposed to secreted ER resident proteins.

KDELRs reside primarily in the Golgi apparatus where they recognize and retrogradely transport proteins with an ERS, or C-terminal “KDEL” or “KDEL-like” sequence, back to the ER lumen^16,17,31,50^. Previous reports from our lab demonstrated that KDELRs interact with the GLuc-SERCaMP construct, and KDELR expression influences ER stress-induced changes in the ER proteome^19,51^. In the present study, we used AAV to overexpress KDELR1 or KDELR2 to examine their ability to protect against EEA-induced exodosis of GLuc-SERCaMP. We found that both KDELR isoforms reduced secretion of GLuc-SERCaMP in response to Tg-or EEA-induced ER Ca^2+^ depletion, which is consistent to a previous report using KDELR1 overexpression in an *in vitro* model of stroke^13^. While there was a similar degree of GLuc-SERCaMP retention by KDELR1 and KDELR2 in the Tg condition, KDELR1 seemed more effective at mitigating glutamate-induced GLuc-SERCaMP secretion vs. KDELR2. This differential effect of KDELR was observed in the 24 h and 30 min glutamate exposure conditions. These differences may be explained by different mechanisms of action of Tg and EEAs to influence ER Ca^2+^ homeostasis. Specifically, Tg induces an irreversible block of the SERCA pump leading to a slow, but steady depletion of ER Ca^2+^. In contrast, glutamate promotes rapid influx of Ca^2+^ from the extracellular space and may induce cellular energy deficits in a manner that is more challenging for the cell to overcome. Additionally, differences in cargo-binding or capacity for cargo between these KDELR isoforms may explain the differences observed in the present study^13,18^.

GLuc-SERCaMP is a robust tool for monitoring the phenomenon of exodosis where ER resident protein secretion occurs following decreases in ER Ca^2+^. In addition to showing that EEAs can induce GLuc-SERCaMP secretion, we also demonstrated that glutamate increased the secretion of the endogenous ER resident protein PDI. In line with the ability of EEA to initiate events at the cell membrane to induce PDI secretion, this effect was mitigated by the NMDAR antagonist, MK801. Further linking the effects of EEAs on PDI redistribution, we demonstrated that overexpression of KDELR blunts the effects of EEA as was observed with GLuc-SERCaMP.

## Conclusions

Herein we provide evidence that EEAs known to cause disruption of intracellular Ca^2+^ homeostasis in models of excitotoxicity also cause a change in ER proteostasis. Our findings expand the current understanding of the molecular consequences of excitotoxicity and support a role for ER Ca^2+^ and ER resident protein secretion in this pathophysiological process. ER stress and altered neuronal Ca^2+^ homeostasis has been documented in humans with temporal lobe epilepsy^52,53^ and in models of seizure^52,54^. Intracellular Ca^2+^ homeostasis impairments have also been documented in models of seizure^40,55–57^. In the present study, we show that GluR activation induces GLuc-SERCaMP secretion, which is indicative of ER Ca^2+^ depletion prompting the redistribution of the ER proteome to the outside of the cell. The effects of EEA-induced GLuc-SERCaMP secretion are carried out, in part, by Ca^2+^ efflux via IP_3_R and RyR. Furthermore, the loss of ER resident proteins in response to EEAs can be overcome in part by increasing the expression of KDELR1 or KDELR2, key regulators of ER protein retention. These findings indicate that a potential therapeutic option for the EEA-induced shift in the ER proteome may include augmenting the KDELR retrieval pathway or by stabilizing ER calcium. Recent work has identified small molecules capable of attenuating ER exodosis triggered by ER calcium depletion which may be of use in disease where excitotoxicity occurs^28^. Our results have implications for understanding the diverse roles of intracellular Ca^2+^ in cell function and survival in conditions of excitotoxicity. Taken together, these data establish a relationship between EEA-induced changes in ER calcium and the secretion of ER resident proteins.

## Acknowledgements

This study was supported by the Intramural Research Program at the National Institute on Drug Abuse. We thank Doug Howard and the NIDA Genetic Engineering and Viral Vector Core for technical contributions.

---

**Extended Data Figure 1-1.**
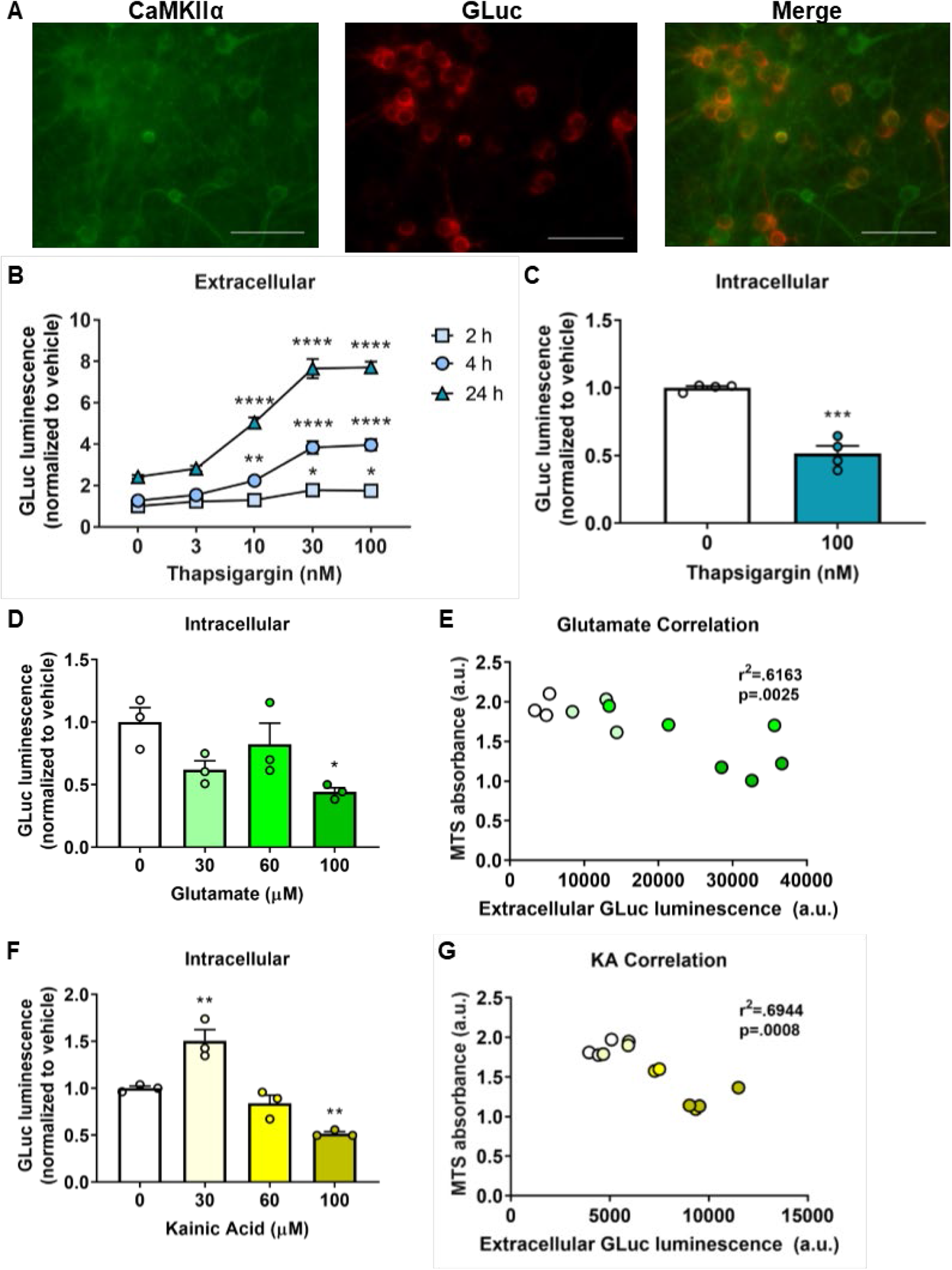
Characterization of AAV1 CaMKIIα GLuc-SERCaMP in rat primary cortical neurons. **A)** Immunofluorescent labeling of endogenous CaMKIIα and the GLuc-SERCaMP in rat primay neurons transduced with AAV1 CaMKIIα GLuc-SERCaMP. **B)** Extracellular (n=5-6/group) and **C,** intracellular levels of GLuc-SERCaMP following treatment with following thapsigargin (Tg) (100 nM) (n=4/group). **D)** Intracellular levels of GLuc-SERCaMP 24 h following treatment with glutamate (n=3/group). **E)** Correlation between GLuc-SERCaMP secretion and viability glutamate treatment (n=3/group, Pearson correlation). **F)** Intracellular levels of GLuc-SERCaMP 24 h following treatment with kainic acid (n=3/group). **G)** Correlation between GLuc-SERCaMP secretion and viability kainic acid treatment (n=3/group, Pearson correlation). Data are shown as mean + SEM. *p<.05, **p<.01 vs. vehicle. *p<.05, ***p<.0005, ****p<.0001 vs. vehicle. 40X magnification, scale bars = 75 µm.

**Extended Data Figure 2-2.**
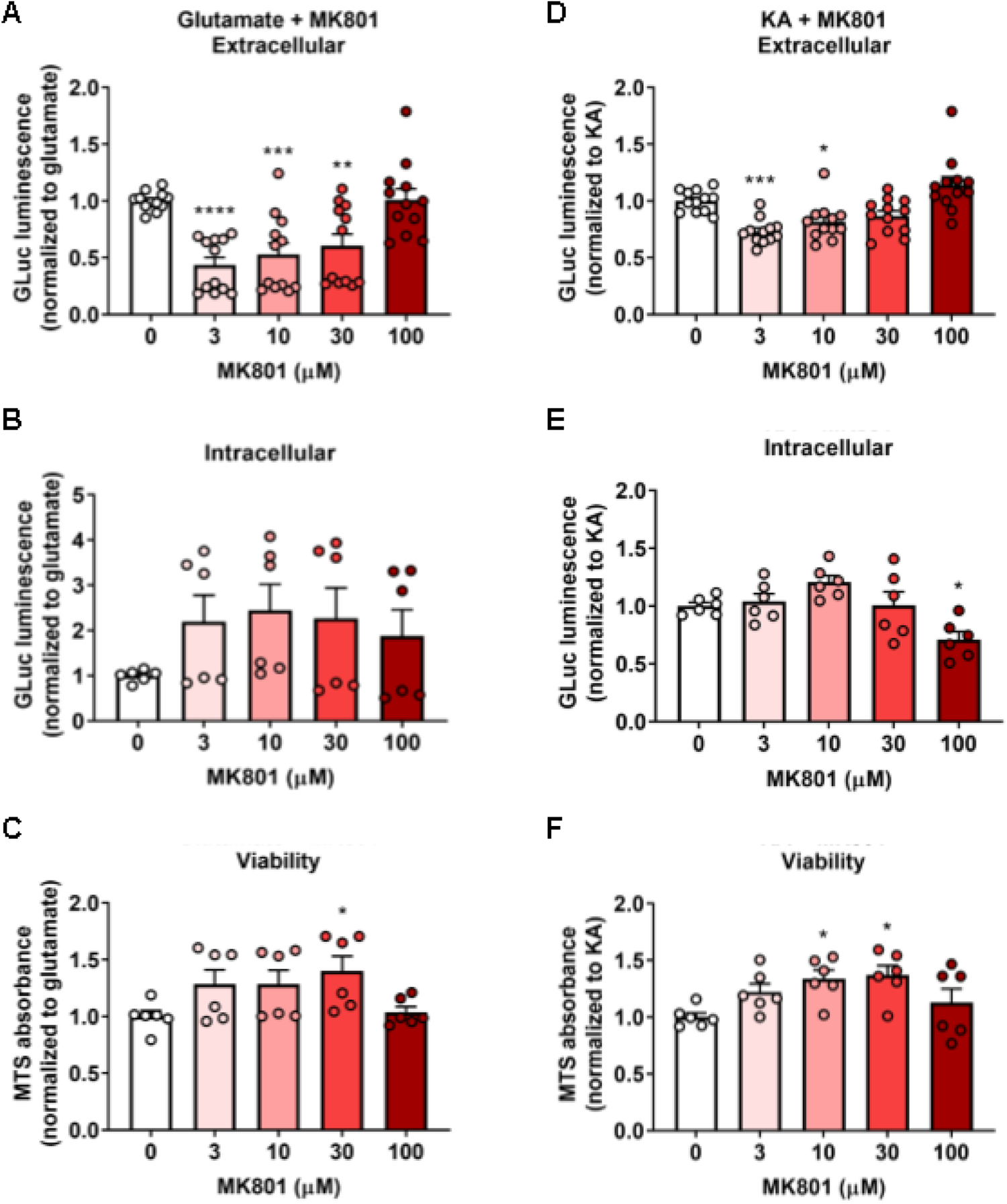
Dose-response evaluation of the NMDA receptor antagonist, MK801, on glutamate and KA-induced changes in ER Ca2+ depletion reporter expression. **A**) MK801 (3, 10, 30, or 100 µM) delivered 30 min prior to glutamate (100 µM), effects on GLuc-SERCaMP secretion (n=12/group). **B**) Effect of MK801 on intracellular levels of GLuc-SERCaMP following treatment with KA (100 µM, n=6/group). **C**) Assay of viability in response to MK801 pre-treatment (n=6/group). **D**) MK801 at 3 and 10 µM blunted the effects of KA to induce GLuc-SERCaMP secretion. E) MK801 did not increase intracellular GLuc-SERCaMP at any dose. **F**) MK801 had modest increase in viability at 10 and 30 µM (**D-E** = 6-12/group). Data are shown as mean + SEM. *p<.05, **p<.01, ***p<.001, ****p<.0001 vs. vehicle.

**Extended Data Figure 3-3.**
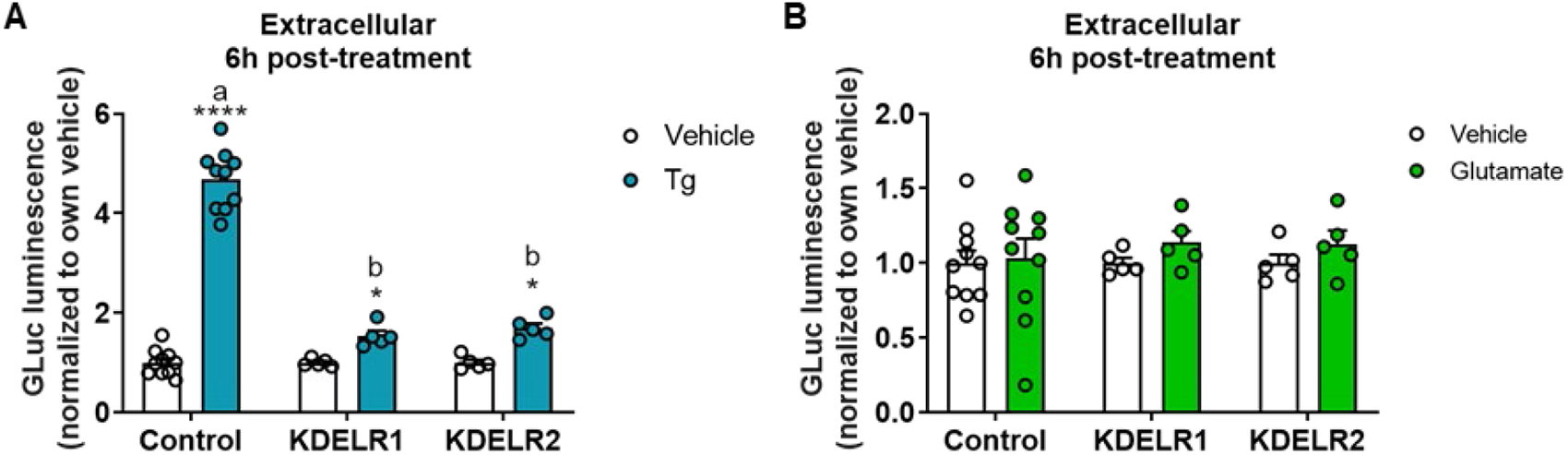
KDELR1 and KDELR2 diminish Tg-induced GLuc-SERCaMP secretion at 6 h post-treatment. **A)** Effect of thapsigargin (100 nM) (n = 5-10/group). **B)** Effect of glutamate (100 µM) on GLuc-SERCaMP secretion in control-, KDELR1-, and KDELR2-expressing cells 6 h post-treatment at 6 h post-treatment (n = 5-10/group. Data are shown as mean + SEM. *p<.05, ****p<.0001 vs. vehicle. Different letters indicate statistically significant differences between AAV conditions, within treatment group.

**Extended Data Figure 5-5.**
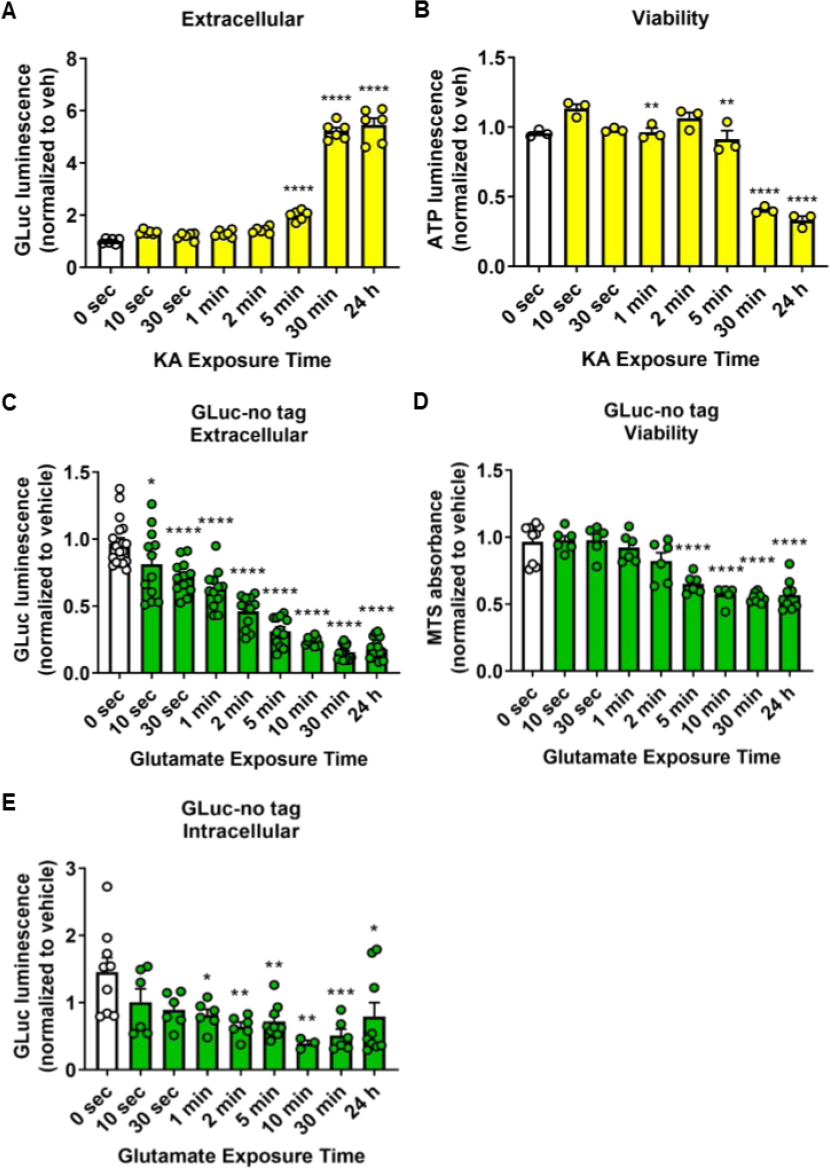
Time-response for KA-induced GLuc-SERCaMP and glutamate-induced GLuc-no tag secretion dynamics. **A)** KA-induced GLuc-SERCaMP secretion (100 µM, n=3/group) and (**B)** viability following ascending durations of exposure (n=3/group). **C)** Glutamate-induced GLuc-no tag secretion (100 µM, n=12-18/group). **D)** Viability of GLuc-no tag-expressing cells following glutamate treatment (100 µM, n=6-9/group). **E)** Intracellular GLuc-no tag levels following glutamate treatment (100 µM, n=3-9/group). Data are shown as mean + SEM, normalized to time-matched vehicle-treated cells.

## Notes

### Competing Interest Statement

The authors have declared no competing interest.

